# Urinary protein changes in the early phase of smoking-induced Chronic Obstructive Pulmonary Disease in a rat model

**DOI:** 10.1101/381053

**Authors:** He Huang, Ting Wang, Youhe Gao

**Affiliations:** Department of Biochemistry and Molecular Biology, Beijing Normal University, Gene Engineering Drug and Biotechnology Beijing Key Laboratory, Beijing, China

**Keywords:** COPD, early urinary biomarkers, proteomics, animal model

## Abstract

Chronic obstructive pulmonary disease (COPD) is a group of severe respiratory diseases. Identifying COPD through early urinary biomarkers by proteomics technology may help to reduce the mortality rate of the disease, improve the quality of life of patients and reduce the burden on society. Urine samples from a COPD rat model induced by smoking were taken at week 2, week 4 and week 8. By LC-MS/MS, 15 differential proteins with human orthologs were identified. After smoking for 2 weeks when there were no significant pathological changes, 8 differential proteins were identified: 2 proteins had been reported to be markers of COPD, while 4 proteins were associated with COPD. After smoking for 4 weeks, which is when slight pathological changes were observed, 7 differential proteins were identified: 3 of them were reported to be associated with COPD, while 1 protein had been reported to be a marker of COPD. After smoking for 8 weeks, there were significant pathological changes: 5 differential proteins were identified, 3 of which were reported to be associated with COPD. The results of this study suggest that differential urinary proteins may provide important clues for the early diagnosis of COPD.

## Introduction

Chronic obstructive pulmonary disease (COPD) is a disease that seriously endangers the breathing of humans, and patients will gradually lose their ability to breathe effectively. It is currently the third leading cause of death in the world [1].

Pulmonary function tests are the gold standard for the clinical diagnosis and assessment of COPD and are the main indicators for determining the airflow limitation. However, in the early stages of COPD, there are usually no airway symptoms and airflow limitation, so pulmonary function tests are difficult to use to diagnose COPD in the early stages of COPD, and in the event of airflow limitation, pulmonary function will be irreversibly damaged[2–5]. We hope to explore new indicators to improve the early diagnosis level of COPD to enable the timely and accurate treatment of patients.

Biomarkers are measurable changes associated with physiological or pathophysiological processes[6]. They are of great significance for the early diagnosis and monitoring of diseases and in the evaluation of treatment effects. Compared with blood, urine lacks mechanisms for maintaining homeostasis and therefore accumulates changes from the body. Through the proteomic analysis of myocarditis[7], glomerulosclerosis[8], ureteral obstruction[9], bacterial meningitis[10], diabetes[11], pulmonary fibrosis[12], subcutaneous tumors[13], and glioblastoma[14] in rat urine, it was found that urine protein can reflect the early changes in various diseases with clinical application prospects.

Cigarette smoke, environmental pollution and bacterial infection are the three dominant risk factors leading to COPD, of which cigarette smoke is the leading cause of COPD[15, 16]. In this study, we establish a cigarette smoke-induced COPD rat model to simulate the pathogenesis of human COPD[17, 18]. By proteomics technology, we look for protein changes associated with COPD in rat urine, hoping to provide a clue for the early diagnosis of COPD.

## Materials and methods

### Animals

Male Wistar’s rats (weight range: 180-200 g; 8 weeks of age) were purchased from Beijing Vital River Laboratory Animal Technology Co., Ltd. After 1 week of conditioning, the rats were randomly divided into a Sham group (clean-air-exposed only n=6) and cigarette-smoke-exposed groups (CS groups n=12). The animal experiments were approved by the Institute of Basic Medical Sciences Animal Ethics Committee, Peking Union Medical College (Approved ID: ACUC-A02-2014-008). All treatments and animal care procedures were performed according to the National Institute of Health guidelines on the care and use of animals.

### Model Establishment

Commercial non-filtered cigarettes (trade name: DA QIAN MEN) containing 11 mg of tar and 0.8 mg of nicotine per cigarette were used in this study. Three rats of model group were kept in a chamber with size of 36 cm (length) × 20 cm (width) × 28 cm (height) and exposed to successive periods of CS at a rate of approximately 10 min per cigarette. At intervals of 1 min, the smoke of a new cigarette was delivered into the chamber and 6 cigarettes for 1 h in the morning and 6 cigarettes 1 h in the afternoon for 6 days a week. After each exposure, the rats were returned to their cages. Control animals (Sham group) inhaled clean (filtered) air only. All of the rats were maintained throughout the study in specific-pathogen-free conditions ventilated with clean air at 20–25 °C. The lights were on a 12-h cycle. At all times, excluding the smoke exposure period, water and diets were provided *ad libitum.*

### Histopathology

The lungs of the rats were removed at week 2, week 4 and week 8 and fixed in 10% neutral-buffered formalin for histopathological analysis. The formalin-fixed tissues were embedded in paraffin, sectioned at 3–5 mm and stained with hematoxylin and eosin (HE) and Alcian blue-periodic acid-Schiff (AB-PAS) to reveal histopathological lesions.

### Urine collection and sample preparation

Urine samples from the COPD rat model induced by smoking were taken at week 2, week 4 and week 8. After collection, the urine was centrifuged at 4 °C for 30 min at 3 000 × g and then at 12 000 × g to remove pellets. Three volumes of ethanol (−20 °C precooling) were added to the supernatant, which was shaken well and then precipitated in a −20 °C refrigerator overnight. The next day, the urine was centrifuged at 4 °C at 12 000 × g for 30 min, and the supernatant was discarded. The pellet was then resuspended in lysis buffer (8 M urea, 2 M thiourea, 50 mM Tris, and 25 mM DTT). The protein concentrations were measured using the Bradford method. Proteins were digested with trypsin (Trypsin Gold, 122 Mass Spec Grade, Promega, Fitchburg,

Wisconsin, USA) using filter-aided sample 123 preparation methods [19]. The peptide mixtures were desalted using Oasis HLB cartridges (Waters, Milford, MA) and dried by vacuum evaporation.

### Liquid chromatography coupled with tandem mass spectrometry (LC-MS/MS) analysis

The digested peptides were acidified with 0.1% formic acid and then loaded onto a reversed-phase micro-capillary column using the Thermo EASY-nLC 1200 HPLC system. The MS data were acquired using the Thermo Orbitrap Fusion Lumos (Thermo Fisher Scientific, Bremen, Germany). The elution gradient for the analytical column was 95% mobile phase A (0.1% formic acid; 99.9% water) to 40% mobile phase B (0.1% formic acid; 89.9% acetonitrile) over 60 min at a flow rate of 300 nL/min. Animals (n=3) with the same clinical manifestations were randomly taken at week 2, week 4, week 8 and from the control group. Each sample was analyzed two times.

### Label-free proteome quantification

The LC-MS/MS results were analyzed using Mascot software and Progenesis software. The database used was the SwissProt_Rat database (551,193 sequences; 196,822,649 residues). The search conditions were trypsin digestion, fixed modification: carbamidomethylation of cysteines, variable modification: oxidation of ethionine, and the tolerances of the parent ion and fragment ion were both 0.05 Da. After normalization, the mass spectrometry peak intensity was used to analyze differential proteins between the control group and COPD group. If the difference between the control group and COPD model group was greater than the intra-group difference, the protein was defined as being a differential protein.

### Gene Ontology Analysis

All of the differentially expressed proteins identified between the control and COPD model groups were analyzed using the OmicsBean database (http://www.omicsbean.cn/) for molecular function, cellular composition, and biological processes.

## Results

### Body weight changes in response to smoke exposure

The control group rats were lively and active with shiny hair, stable breathing and rapid weight gain. The cigarette smoke group rats look tired, and after 1 week of cigarette smoke, their weight was significantly lower than the weight of those in the control group (*: P <0.01).

### Histological changes

In the lung parenchyma, there were no significant changes in the bronchus (Fig. 4B) and alveoli (Fig. 5B) in the week 2 group. Four weeks after smoking, bronchial epithelial detachment was observed (Fig. 4C) and the alveolar space expanded and ruptured (Fig. 5C). At week 8, the rat lung bronchial epithelial cells were denatured, adhered, and partially detached (Fig. 4D) and the alveolar wall became thinner, while the alveolar space expanded, ruptured or had bullae formed in it (Fig. 5D). Compared with the control group, AB-PAS staining showed no significant change in the week 2 group (Fig. 6B). At week 4, bronchial epithelial goblet cells in the rats became larger (Fig. 6C). At week 8, the number of goblet cells (in blue) in the bronchial epithelium was dramatically elevated and the size of goblet cells was enlarged with hypertrophy and hypersecretion (Fig. 6D).

**Figure 1.**
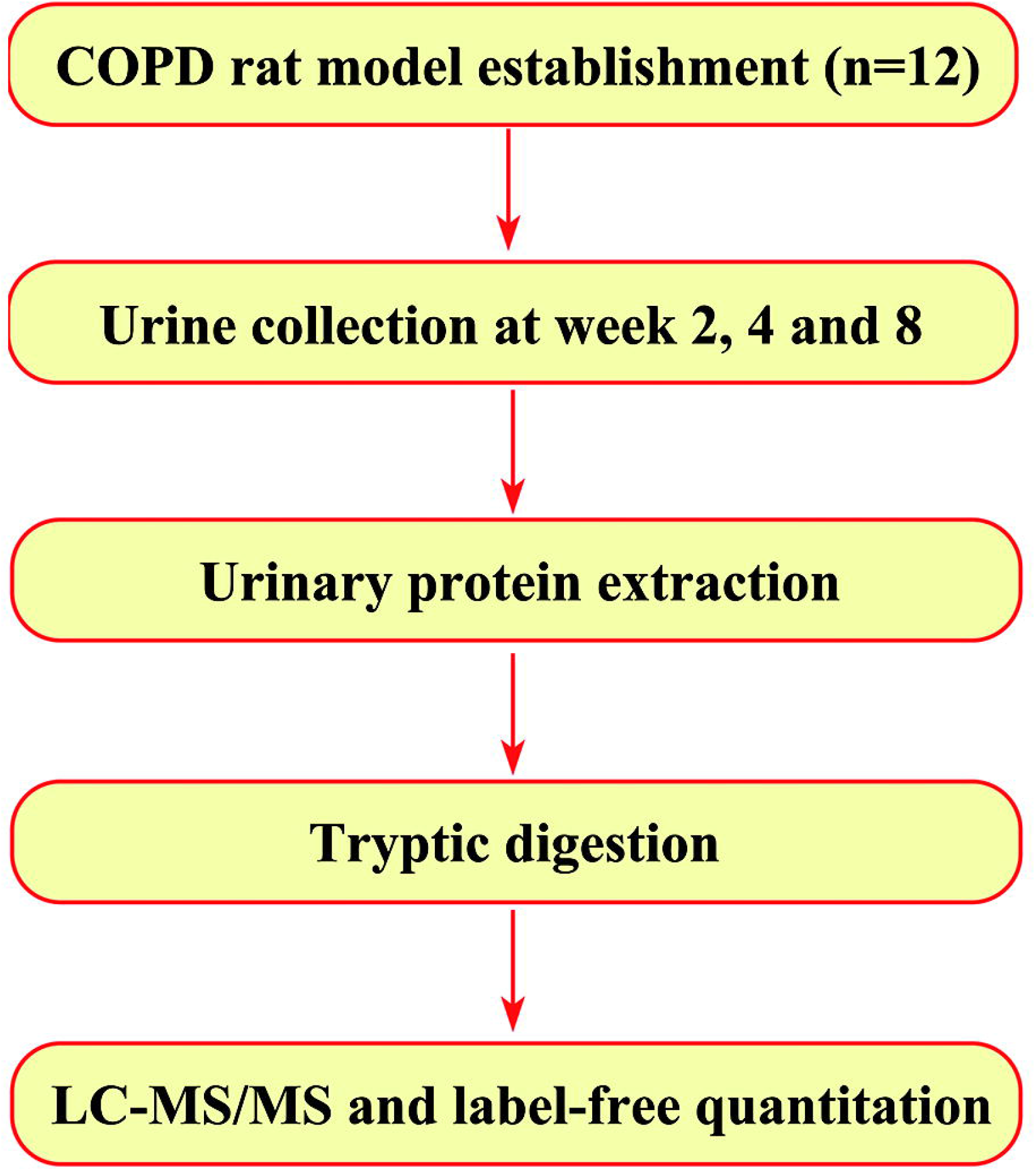
Workflow of urinary protein changes analysis in this study. Urine samples from a COPD rat model induced by smoking were taken at week 2, week 4 and week 8, and the urinary proteome was analyzed using liquid chromatography coupled with tandem mass spectrometry (LC-MS/MS) identification.

**Figure 2.**
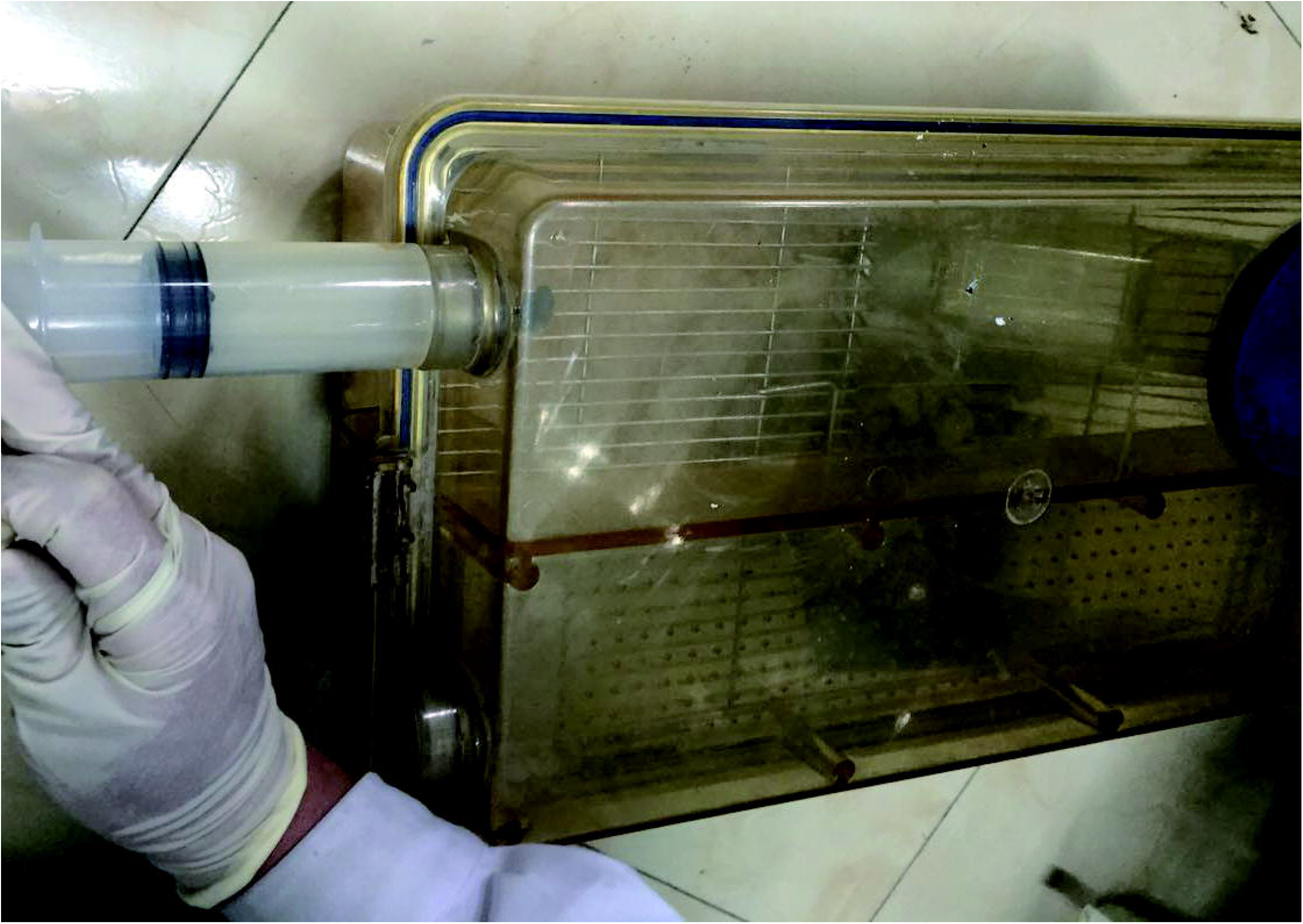
Establishment of COPD rat model. Three rats of model group were kept in a chamber with size of 36 cm (length) × 20 cm (width) × 28 cm (height) and exposed to successive periods of CS at a rate of approximately 10 min per cigarette. At intervals of 1 min, the smoke of a new cigarette was delivered into the chamber and 6 cigarettes for 1 h in the morning and 6 cigarettes 1 h in the afternoon for 6 days a week. After each exposure, the rats were returned to their cages. Control animals (Sham group) inhaled clean (filtered) air only.

**Figure 3.**
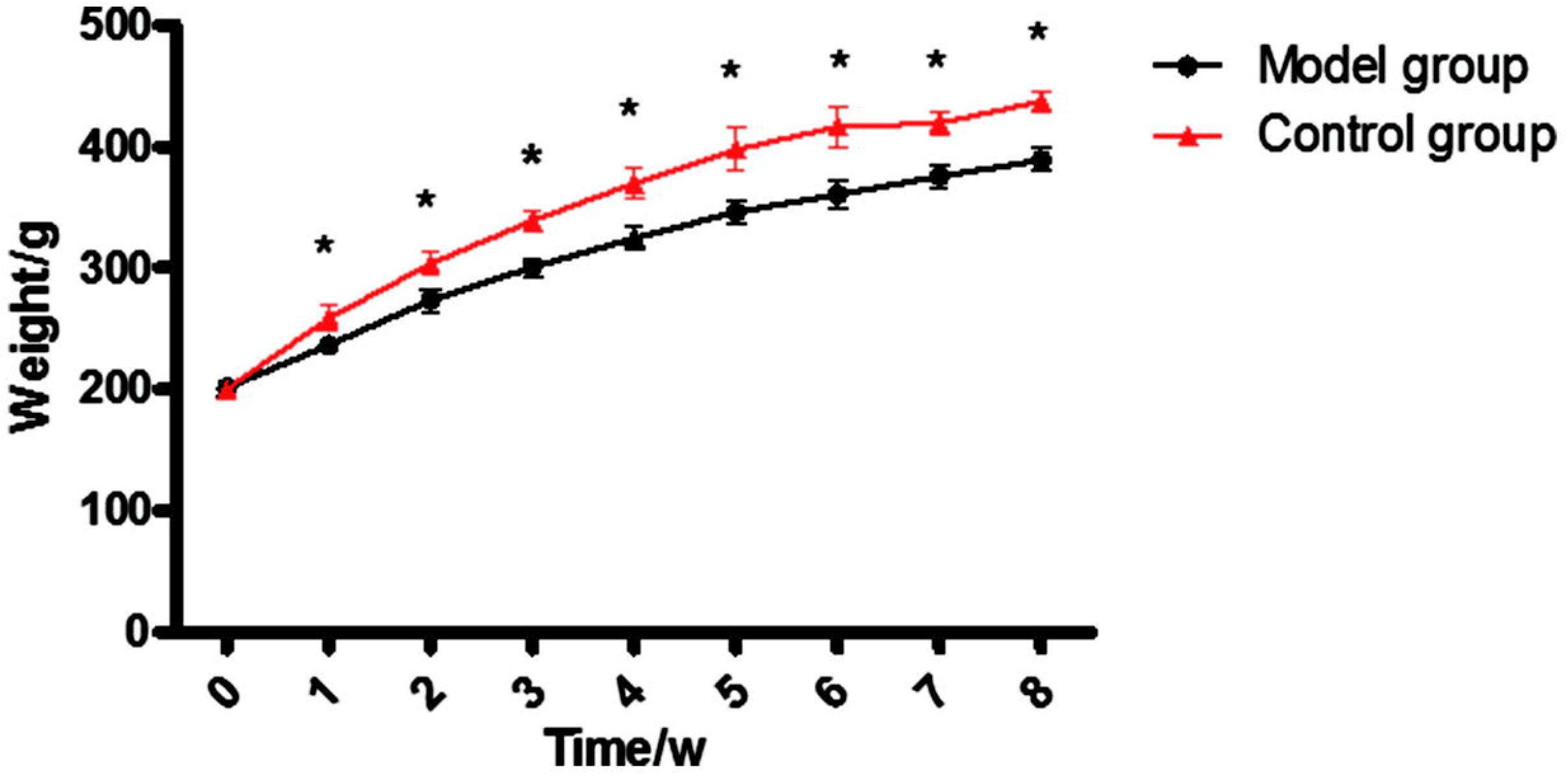
Time course for the body weight of the COPD rat model. Since being exposed to cigarette smoke for 1 week, the body weight of the COPD group was significantly lower than that of the control group. The values shown are the mean±S.E.M for n=8-12 rats in the COPD group and n=3-6 rats in the control group; * indicates p < 0.01.

**Figure 4.**
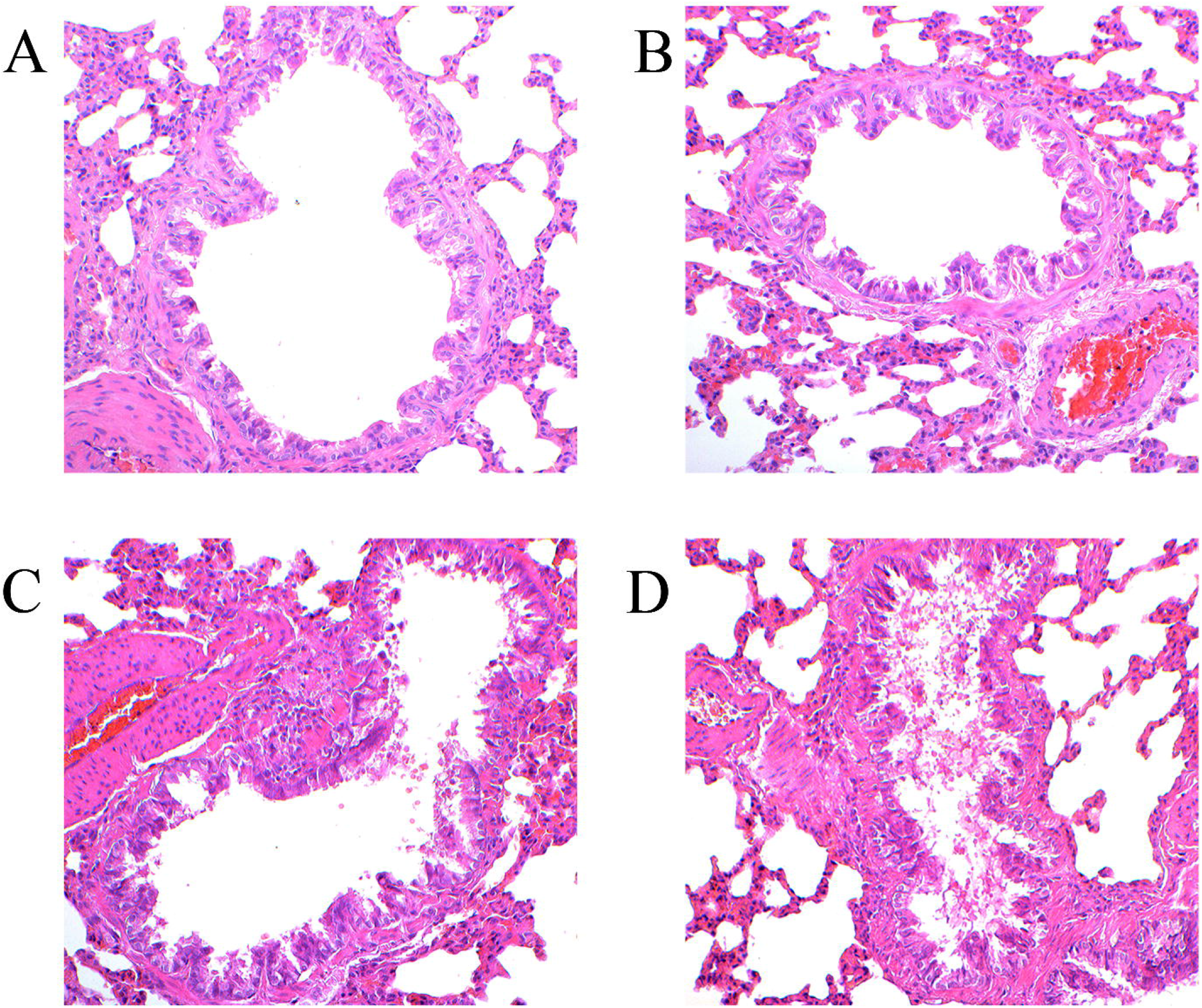
Histological examination of the rat bronchial tissues after inhalation of cigarette smoke (CS). Sections of bronchial tissues were stained with hematoxylin–eosin and photographed under a microscope. (A) Control group, (B) CS exposure for 2 wk, (C) CS exposure for 4 wk, and (D) CS exposure for 8 wk. Original magnification: 200×.

**Figure 5.**
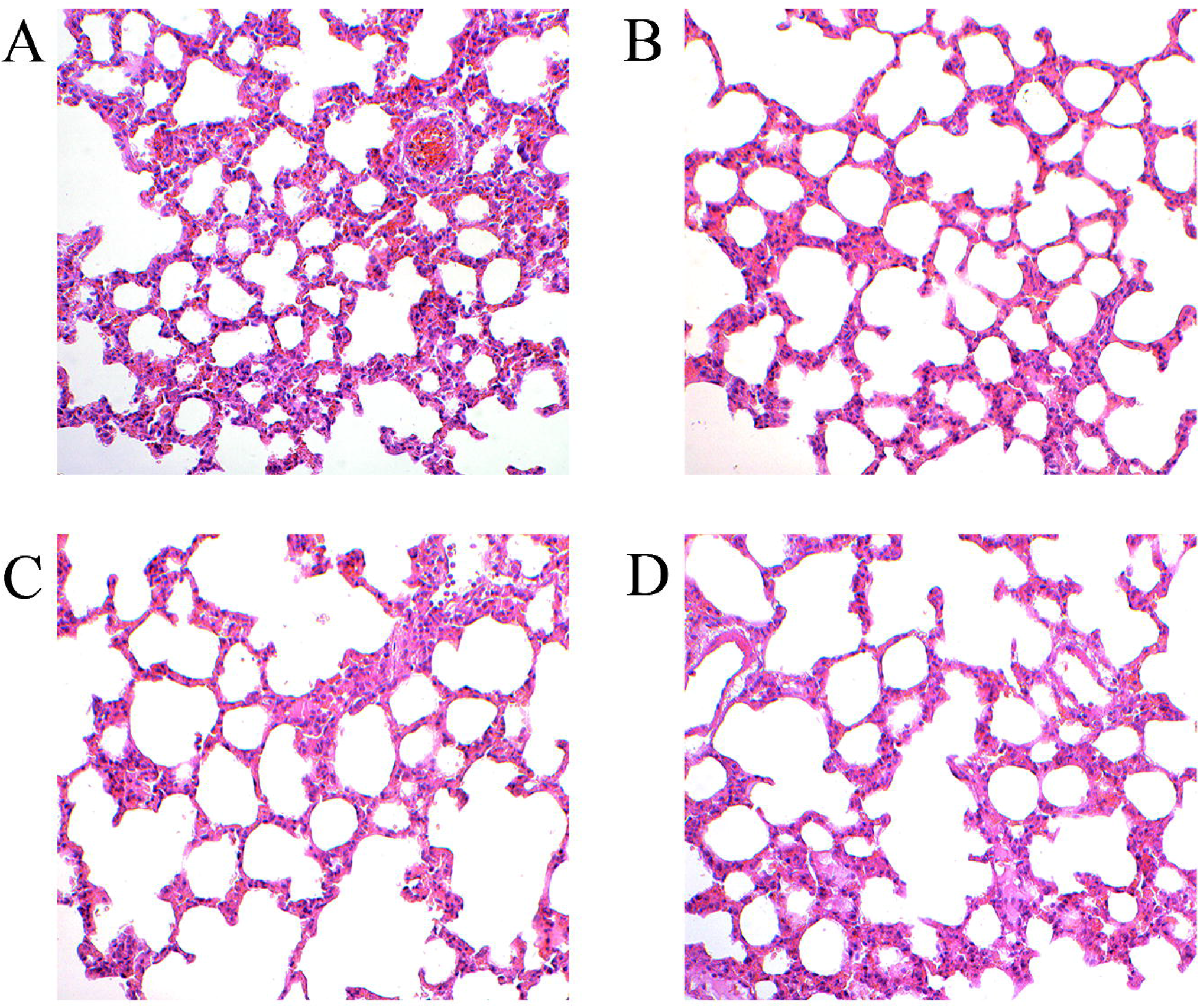
Histological examination of the rat lung tissues after inhalation of cigarette smoke (CS). Sections of lung parenchyma were stained with hematoxylin–eosin and photographed under a microscope. (A) Control group, (B) CS exposure for 2 wk, (C) CS exposure for 4 wk, and (D) CS exposure for 8 wk. Original magnification: 200×.

**Figure 6.**
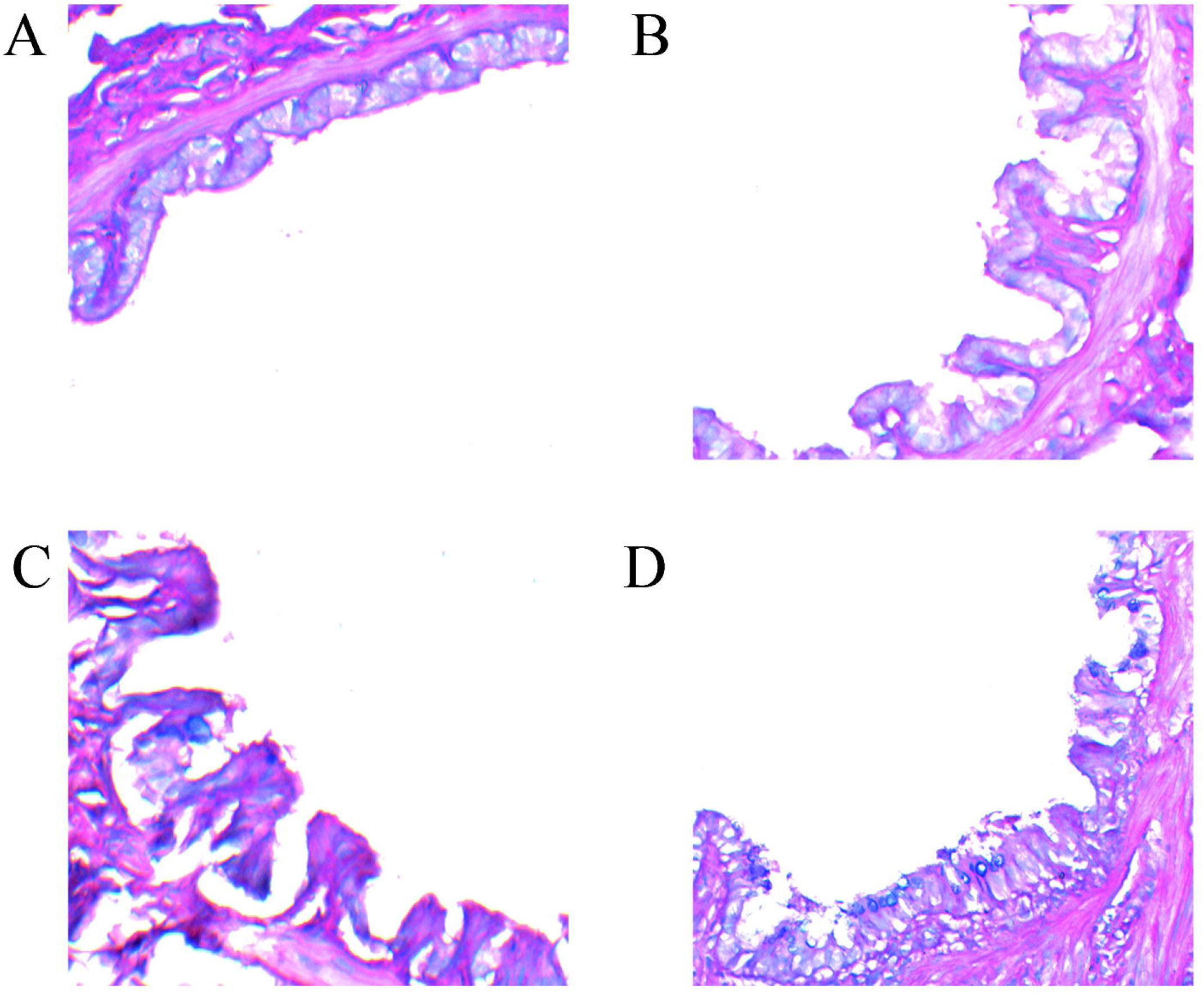
Alcian blue-periodic acid-Schiff (AB-PAS) staining for mucus expression in the epithelium of the rat bronchus after inhalation of cigarette smoke (CS). (A) Control group, (B) CS exposure for 2 wk, (C) CS exposure for 4 wk, and (D) CS exposure for 8 wk. Original magnification: 200×.

### Differential urinary proteins

In total, 547 urine proteins were identified, and 21 differential proteins were selected according to the following criteria: 1) proteins with at least two specific peptides were identified, 2) a fold change> 1.5; and 3) a p value < 0.05. Among the differential proteins, 15 had corresponding human orthologs. There were 8 significantly differentially expressed proteins (Table 1) at week 2, 7 significantly changed proteins at week 4 (Table 2), and 5 significantly changed proteins at week 8 (Table 3). The common proteins between the week 2 group and week 4 group were fetuin-A, plasminogen and galectin-3-binding protein. Nucleoside diphosphate kinase B and phosphoglycerate kinase-1 were the common proteins between the week 4 and week 8 groups.

**Table 1.** Differentially expressed proteins in the urine of rats with COPD at week 2. 8 differential proteins were identified: 2 proteins had been reported to be markers of COPD, while 4 proteins were associated with COPD.

| Accession | Human | Protein description | Trend | Fold change | COPD Correlation | COPD Biomarker |
| --- | --- | --- | --- | --- | --- | --- |
| P02651 | P06727 | Apolipoprotein A-IV | ↑ | 4.04 |  |  |
| P04276 | P02774 | Vitamin D-binding protein | ↑ | 2.54 | [21] | [20,22,23,24] |
| Q01177 | P00747 | Plasminogen | ↑ | 2.37 | [27] |  |
| P24090 | P02765 | Fetuin-A | ↑ | 2.22 | [26] | [25] |
| Q05175 | P80723 | Brain acid soluble protein-1 | ↑ | 1.99 |  |  |
| Q63467 | P04155 | Trefoil factor-1 | ↑ | 1.86 | [28] |  |
| Q5XI43 | Q9BRK3 | Matrix remodeling-associated protein-8 | ↑ | 1.63 |  |  |
| O70513 | Q08380 | Galectin-3-binding protein | ↓ | 0.44 |  |  |

**Table 2.** Differentially expressed proteins in the urine of rats with COPD at week 4. 7 differential proteins were identified: 3 of them were reported to be associated with COPD, while 1 protein had been reported to be a marker of COPD.

| Accession | Human | Protein description | Trend | Fold change | COPD Correlation | COPD biomarker |
| --- | --- | --- | --- | --- | --- | --- |
| P24090 | P02765 | Fetuin-A | ↑ | 2.94 | [26] | [25] |
| P19804 | P22392 | Nucleoside diphosphate kinase B | ↑ | 2.86 | [31] |  |
| P42123 | P07195 | L-lactate dehydrogenase B chain | ↑ | 2.67 |  |  |
| Q01177 | P00747 | Plasminogen | ↑ | 2.27 | [27] |  |
| P16617 | P00558 | Phosphoglycerate kinase-1 | ↑ | 1.62 |  |  |
| Q9JJ40 | Q5T2W1 | Na(+)/H(+) exchange regulatory cofactor-3 | ↑ | 1.51 |  |  |
| O70513 | Q08380 | Galectin-3-binding protein | ↓ | 0.35 |  |  |

**Table 3.** Differentially expressed proteins in the urine of rats with COPD at week 8. 5 differential proteins were identified, 3 of which were reported to be associated with COPD.

| Accession | Human | Protein description | Trend | Fold change | COPD Correlation | COPD Biomarker |
| --- | --- | --- | --- | --- | --- | --- |
| P19804 | P22392 | Nucleoside diphosphate kinase B | ↑ | 7.24 | [31] |  |
| O88989 | P40925 | Malate dehydrogenase, cytoplasmic | ↑ | 2.32 | [33] |  |
| P16617 | P00558 | Phosphoglycerate kinase-1 | ↑ | 2.24 |  |  |
| P00758 | P06870 | Kallikrein-1 | ↑ | 2.11 | [34] |  |
| P07314 | P19440 | γ-glutamyltranspeptidase K89 | ↑ | 2.05 |  |  |

The expression of vitamin D-binding protein (VDBP) increased at week 2. James D. Bortner[20] found that VDBP expression was significantly down-regulated in smoker plasma and could be used as a candidate marker of smoking-induced disease. A.M. Wood [21]reported that VDBP relates to FEV1 and affects macrophage activation. In addition, there is a close association between COPD and VDBP gene polymorphisms and the VDBP gene polymorphism could be a potential marker used for the screening of COPD[22–24]. The study suggests that fetuin-A is a reproducible and clinically relevant biomarker in patients with COPD that may be useful in the identification of exacerbation-prone patients[25] and that serum fetuin-A levels are associated with carotid intima–media thickness in patients with normotensive chronic obstructive pulmonary disease[26]. Plasminogen is not reported to be directly related to COPD, but plasminogen activator inhibitor-1 may be a potential biomarker candidate for COPD-specific and extra-pulmonary manifestation studies [27]. As for trefoil factor-1, there is an increased secretion of trefoil factor (TFF) peptides in the lungs of patients with COPD, as well as significant increases in their serum levels, which suggests a role of TFF peptides in the mucus hypersecretion in pulmonary diseases[28]. Brain acid-soluble protein-1 was found to be up-regulated in lung adenocarcinoma A549 cells[29]. Galectin-3-binding protein was selected as one of the candidate proteins for specifically diagnosing large-cell neuroendocrine lung carcinoma (LCNEC) and differentiating it from other lung cancer subtypes[30]. Apolipoprotein A-IV and matrix remodeling-associated protein-8 are newly discovered urinary proteins associated with COPD in this study.

At week 4, fetuin-A and plasminogen still showed higher expression levels, while galectin-3-binding protein is still down-regulated. Cystic fibrosis results from mutations in the cystic fibrosis transmembrane conductance regulator (CFTR) nucleoside diphosphate kinase B, which forms a functional complex with CFTR [31]. Additionally, phosphoglycerate kinase-1 (PGK1) and metaLnc9 facilitate lung cancer metastasis via a PGK1-activated AKT/mTOR pathway[32]. The L-lactate dehydrogenase B chain and Na(+)/H(+) exchange regulator-3 are newly discovered urinary proteins associated with COPD in this study.

At week 8, nucleoside diphosphate kinase B and phosphoglycerate kinase-1 were still significantly up-regulated. The expression level of malate dehydrogenase in the lung of asthmatic mice can help assess lung mitochondrial damage, which can lead to respiratory diseases[33]. KLK1 (tissue kallikrein 1) is a member of the tissue kallikrein family of serine proteases and is the primary kinin-generating enzyme in human airways [34]. For the first time, γ-glutamyltranspeptidase K89 was found to be associated with COPD in this study.

### Functional analysis of differential proteins

GO analysis of the differential proteins in the urine of the COPD rats by OmicsBean was performed. The differential proteins were mainly divided into three categories: transport proteins, binding proteins and enzyme active proteins. The biological process analysis showed that the differential proteins in the week 2 group were mainly involved in the processes of wound response and homeostasis (Fig. 7A). Most of the differential proteins showed apolipoprotein binding, phosphatidylcholine binding, cholesterol transport, sterol transport and a cholesterol-binding molecular function. Compared with those in the week 2 group, the differential proteins in the week 4 group (Fig. 7B) and week 8 group (Fig. 7C) were mainly involved in the metabolism of nucleotides, pyridine-containing compounds, and small base-containing molecules. The week 4 group of differentially expressed proteins showed molecular function of scavenger receptor binding and found that the scavenger receptor induced relevant signaling pathways of innate immune responses to smoke-induced COPD[35]. In addition, one main characteristic of COPD is the thickening of the airway smooth muscle, which is partly due to airway smooth muscle cell (ASMC) hyperplasia. Additionally, changes in glycolysis, glutamine and fatty acid metabolism may lead to increased biosynthesis and redox balance, supporting COPD ASMC growth[36]. This correlates with the GO analysis results of the differential proteins of the week 8 group in this study. Moreover, most of these differentiated proteins originate from extracellular components. Some proteins at week 8 are derived from mitochondria. One study[37] shows that the mitochondria could be a crucial contributor to immunity and that mitochondrial dysfunction often has a direct impact on health, such as lung injury and COPD.

**Figure 7.**
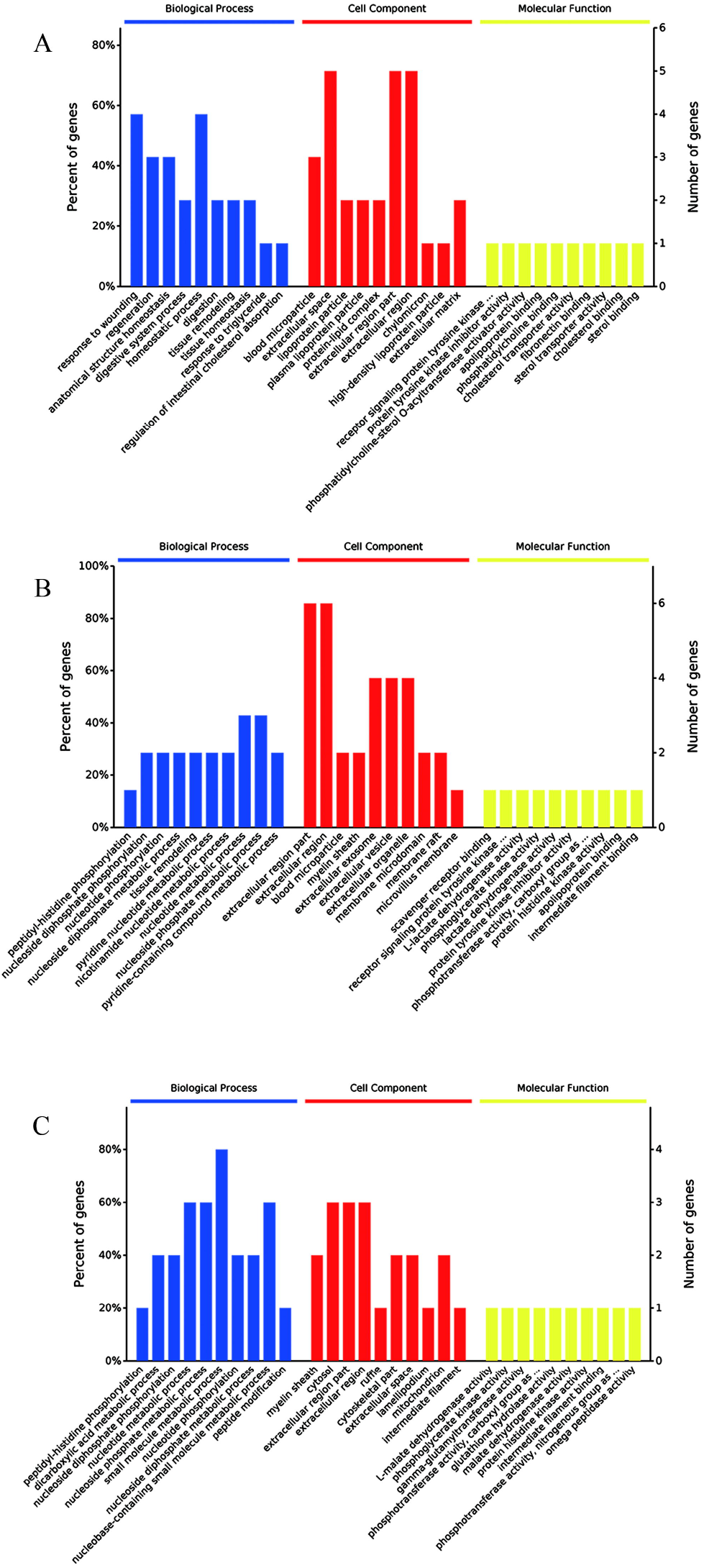
GO analysis of the differential proteins. (A) week 2 group, (B) week 4 group, and (C) week 8 group.

## Discussion

In this study, we established a COPD rat model using cigarette smoke to simulate the pathogenesis of COPD and monitored the very early protein changes in the urine of COPD rats. Moreover, variables that have an impact on clinical samples, such as medication, surgery, and patients’ living habits and so on, were excluded. Helping to reveal the pathogenesis of COPD and improve the diagnostic level of COPD is of great significance. As a preliminary study, we found that urine has great promise in finding early biomarkers associated with COPD. In future studies, a large number of clinical urine samples from COPD patients are needed to verify the protein patterns of specific biomarkers. This will enhance our understanding of COPD and help us better understand the role of urine in the search for biomarkers of respiratory diseases.

## Conclusion

In this study, some markers that have been reported to be associated with COPD can be detected in the very early phase in COPD rat model urine.

## Author Contributions

H.H. and Y.G. prepared the first draft. H.H, T.W. and Y.G. conceived and designed the experiments. H.H. and T.W. performed the experiments and analyzed the data. All authors approved the final manuscript.

## Acknowledgements

This work was supported by the National Key Research and Development Program of China (2016 YFC 1306300), Key Basic Research Program of the Ministry of Science and Technology of China (2013FY114100), Beijing Natural Science Foundation (Nos. 7173264 and 7172076), Beijing cooperative construction project (110651103) Beijing Normal University (11100704) Peking Union Medical College Hospital (2016-2.27).

---

**Supplemental File 1.** All identified urinary proteins and the details in this study.

